# Morphological study of embryonic *Chd8^+/-^* mouse brains using light-sheet microscopy

**DOI:** 10.1101/2020.10.05.326132

**Authors:** Harold F. Gómez, Leonie Hodel, Odyssé Michos, Dagmar Iber

## Abstract

**Objective:** Autism spectrum disorder (ASD) encompasses a group of neurodevelopmental conditions that remain poorly understood due to their genetic complexity. *CHD8* is a risk allele strongly associated with ASD, and heterozygous *Chd8* loss-of-function mice have been reported to exhibit macrocephaly in early postnatal stages. In this work, we sought to identify measurable brain alterations in early embryonic development.

**Results:** We performed light-sheet fluorescence microscopy imaging of N-cadherin stained and optically cleared *Chd8*^*+/-*^ and wild-type mouse brains at embryonic day 12.5 (E12.5). We report a detailed morphometric characterization of embryonic brain shapes and cortical neuroepithelial apical architecture. While *Chd8*^*+/-*^ characteristic expansion of the forebrain and midbrain was not observed this early in embryogenesis, a tendency for a decreased lateral ventricular sphericity and an increased intraocular distance in *Chd8*^*+/-*^ brains was found compared to controls. This study advocates the use of high-resolution microscopy technologies and multi-scale morphometric analyses of target brain regions to explore the etiology and cellular basis of *Chd8* haploinsufficiency.

## Introduction

Autism spectrum disorder (ASD) is a poorly understood disease due to significant genetic complexity and phenotypic heterogeneity. Despite our improved understanding of how ASD develops (1–5), mapping the relative contribution of risk alleles to neuroanatomical abnormalities and clinically observed phenotypes like macrocephaly remains challenging.

Genetic studies have implicated mutations in more than 800 genes of diverse function in the etiology of ASD (1). One of the most strongly associated genes is the chromodomain helicase DNA-binding protein 8 *CHD8*, an ATP-dependent chromatin remodeler, and transcriptional repressor. Patients with loss-of-function (LOF) mutations in *CHD8* exhibit gene haploinsufficiency, display altered behavior, and region-specific anomalies in brain morphology and physiology that manifest during early childhood (6–8). Similarly, *Chd8* haploinsufficiency in mice results in neonatal macrocephaly, increased brain weight, and craniofacial abnormalities (9–11), mirroring clinical observations in patients and suggesting similar developmental trajectories between species.

ASDs are largely hypothesized to originate in utero from profound perturbations in neural stem cell niche regions of the developing brain (12). Gene expression profiling in the embryonic mouse cortex of *Chd8* haploinsufficient mice shows a temporal modulation of *Chd8* that peaks at E12, and helps negotiate the complex balance between neuronal expansion (prior to E12.5) and differentiation (E12.5 to postnatal) (13). As a result, dysregulations of *Chd8* dynamics during embryonic cortical development in mice prematurely deplete the neural progenitor pool, consistent with a lower density of neural cells and metabolic components observed in children with ASD (14). In spite of novel neurodevelopmental evidence, it is unknown whether these in utero perturbations manifest as distinctive anatomical dysmorphologies before the postnatal onset of characteristic ASD phenotypes (15).

In this study, we investigated the morphological consequences of *Chd8* haploinsufficiency during embryonic mouse brains at the whole-organ and cellular level. To anticipate anatomical findings in a condition with early-life onset, we leveraged N-cadherin staining, light-sheet fluorescence microscopy, and CUBIC tissue clearing (16) to examine the neuroanatomical differences between E12.5 mouse brains with germline *Chd8*^+/-^ LOF mutations (10) and litter-matched wild-types. We report slight differences in intraocular distance and ventricular sphericity and introduce a detailed approach for comparing cortical neuroepithelial apical architecture. Taken together, these datasets provide a new avenue for querying the developmental role of *CHD8* and the cellular remodeling that is likely to precede associated post-birth brain malformations in haploinsufficiency cases.

## Methods and materials

### Animals

Mice with loss-of-function mutations in *Chd8* were generated using Cas9-mediated germline editing (10).

### Immunofluorescence on embryonic brains

Following dissection, all E12.5 mouse brains were fixed with 4% paraformaldehyde in PBS. Samples were then incubated with anti-N-cadherin antibody (BD Transduction Laboratories; Material No. 610920; 1:200) at 4 °C for 3 days. After washing in D-PBS, brains were incubated with conjugated fluorescent secondary Alexa Fluor 555 donkey anti-mouse IgG (H+L) (Abcam; Material No. ab150106; 1:250) for 2 days at 4 °C.

### Optical clearing and light-sheet imaging

Whole-mount clearing was performed with the Clear Unobstructed Brain/Body Imaging Cocktails and Computational Analysis (CUBIC) protocol (16). Delipidation and refractive index matching were carried out with reagent-1 [25% (w/w) urea, 25% ethylenediamine, 15% (w/w) Triton X-100 in distilled water] and reagent-2 [25% (w/w) urea, 50% (w/w) sucrose, 10% (w/w) nitrilotriethanol in distilled water], respectively. Samples were incubated in 1/2 reagent-1 (CUBIC-1:H2O=1:1) for 1 day and then in 1X reagent-1 until transparent. All samples were washed several times in PBS and treated with 1/2 reagent-2 (CUBIC-2:PBS=1:1) for around 3 days. Lastly, incubation in 1X reagent-2 was done until transparency was achieved, and the solution became homogeneous. All steps were performed on a shaker at room temperature.

Fluorescence images were acquired using a Zeiss Lightsheet Z.1 microscope. Acquisition optics included a Zeiss 20x/1.0 Plan Apochromat water-immersion objective to acquire cell resolution data, and a Zeiss 5x/0.1 air objective lens for larger fields of view (whole brains). All image stacks were deconvolved using Huygens deconvolution to improve contrast and resolution and further pre-processed in Fiji (17) to accentuate feature boundaries.

### 3D surface reconstruction of whole mouse brains

3D segmentation of the embryonic ventricles and cerebral cortex was conducted with Imaris MeasurementPro, a component of Imaris v9.1.2 (BitPlane, South Windsor, CT, USA). This enabled the computational interpolation of planar 2D surface outlines from successive horizontal sections into 3D iso-surface. Contours were drawn on magnified images to allow sub-voxel precision and faithful delineation of small-scale features (Fig. 1). Quantified brain surface features included volume and surface area. Intraocular distance was measured in 3D using measurement points placed in the center of the pupils.

**Figure 1.**
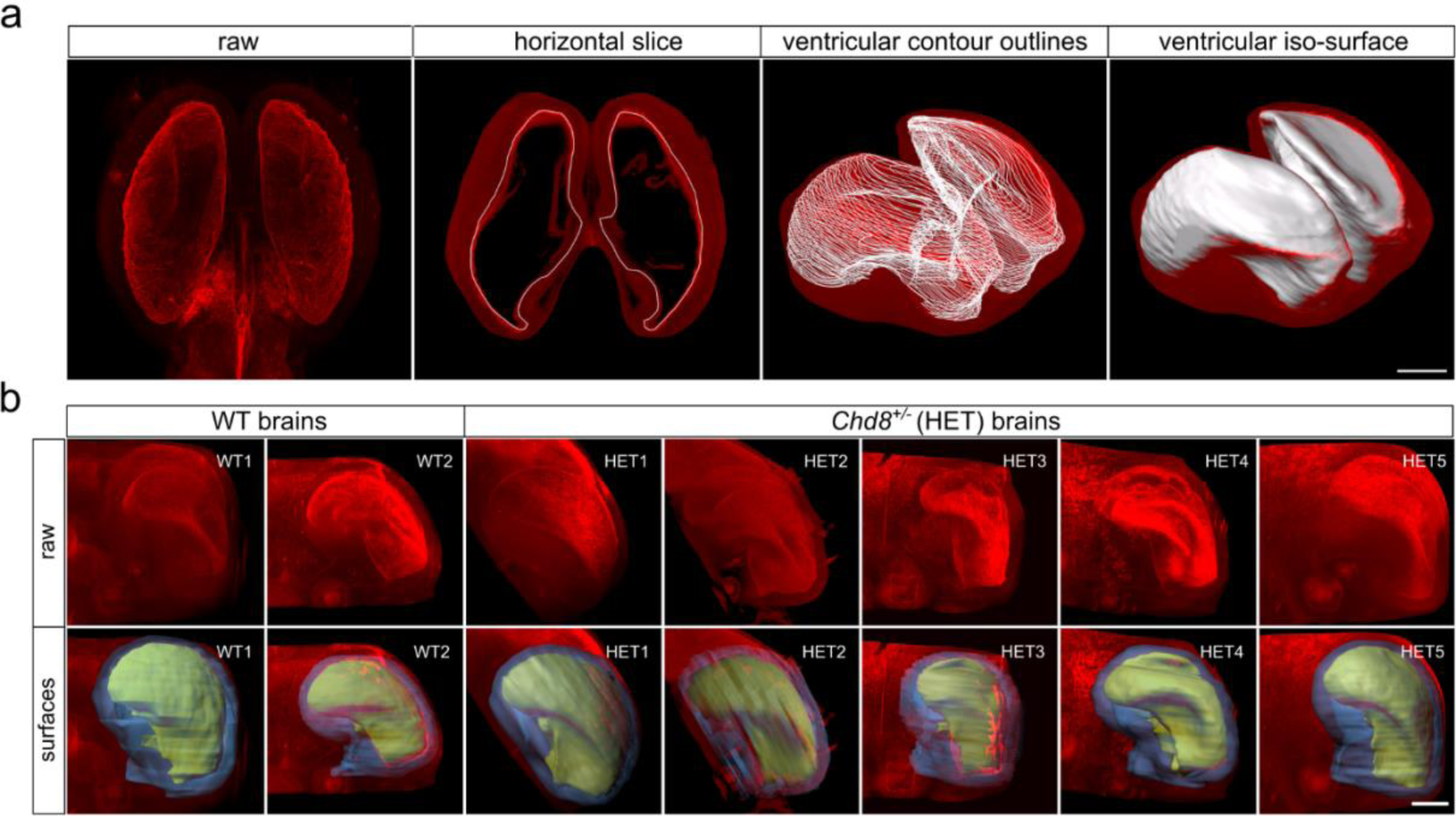
Volumetric analysis of CUBIC-cleared wild-type and *Chd8*^*+/–*^ samples, immunostained for N-cadherin (red) to mark neuronal epithelial tissue. **(a)** Illustration of the processing steps in the creation of manual surfaces. Sequential ventricular contours were drawn manually throughout the entire dataset to extract 3D morphology (white outlines). Scale bar 500 µm. **(b)** Raw (top row) and overlays (bottom row) of ventricular (yellow) and cortical (blue) iso-surfaces for each sample. Scale bar 400 µm.

As the cortex has broad irregular anatomical features that make absolute cortical thickness measurements challenging, we considered the cortex to be a hollow cylinder with volume *V* and area *A*. In this way, the cortical height of the neuroepithelial layer could be approximated as

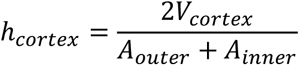

Furthermore, to derive ventricular sphericity, lateral ventricle iso-surfaces were separated at the septum pellucidum. The sphericity of each 3D entity was then determined as

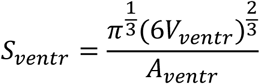

### Morphometric measurements of apical neuroepithelia

Cell morphology in the apical layer of the cortical epidermis was investigated using the open-source software platform MorphoGraphX (18). A curved 2.5D image projection was constructed by meshing the apical boundary and projecting 2-6 µm of the most apical signal onto it. Then, the Watershed algorithm was used to segment all cell boundaries, some of which required manual curation. All border cells were excluded from the analysis.

To characterize all projected polygonal apical lattices, quantifications on cellular areas and neighbour numbers were imported into the R software platform. Apical packing was also explored via known regularities of epithelial lattices. Termed Lewis’ Law, this property linearly relates the measured average cell area *Ā* and neighbour number *n* and has been previously described in all apical epithelia studied to date (19,20).

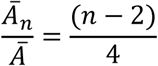

As cells with small polygon numbers have the tendency to be in contact with cells of larger polygon numbers and vice-versa, one also observes that the average number of neighbours of all *n* cells that border a cell with *n* neighbours follows

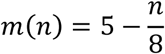

a relationship termed Aboav-Weaire’s Law (21,22). Lastly, the cell aspect ratio was calculated using an in-house algorithm that leverages MorphoGraphX’s modularity to fit an ellipse and extract major and minor axes for each cell outline.

## Results

### ASD-associated craniofacial phenotypes in Chd8^+/-^ mice

To determine whether *Chd8* heterozygous mice exhibit structural and craniofacial ASD phenotypes during embryonic development, we tested for differences in brain morphology between E12.5 *Chd8*^*+/-*^ and control animals. To this extent, user-assisted 3D segmentation software tools were used to derive iso-surface representations from volumetric image stacks and enable tissue quantification measures of size, shape, and asymmetry (Fig. 1).

We then characterized different anatomical features in both cortical and ventricular regions to reveal regional alterations (Fig. 2). Overall brain volume, including ventricles, showed no significant difference between the two groups (wild-type [n=2] 3.41 mm^3^ and 3.15 mm^3^, *Chd8*^+/-^ [n=5] 3.60 ± 0.56 mm^3^) (Fig. 2a). Similarly, measured cortical volumes excluding the ventricular space showed no difference (wild-type [n=2] 2.17 mm^3^ and 2.15 mm^3^, *Chd8*^*+/-*^ [n=5]: 2.47 ± 0.37 mm^3^). Individual ventricular volume was consistent within and between groups (wild-type [n=4] 0.56 ± 0.07 mm^3^, *Chd8*^*+/-*^ [n=10] 0.57 ± 0.14 mm^3^). Furthermore, we observed no differences in brain surface area (wild-type [n=2] 15.6 mm^2^ and 14.5 mm^2^, *Chd8*^*+/-*^ [n=5] 15.48 ± 1.63 mm^2^) or in ventricular surface area (wild-type [n=4] 4.66 ± 0.30 mm^2^, *Chd8*^*+/-*^ [n=10] 4.95 ± 0.83 mm^2^) between *Chd8*^*+/-*^ and wild-type littermates (Fig. 2b).

**Figure 2.**
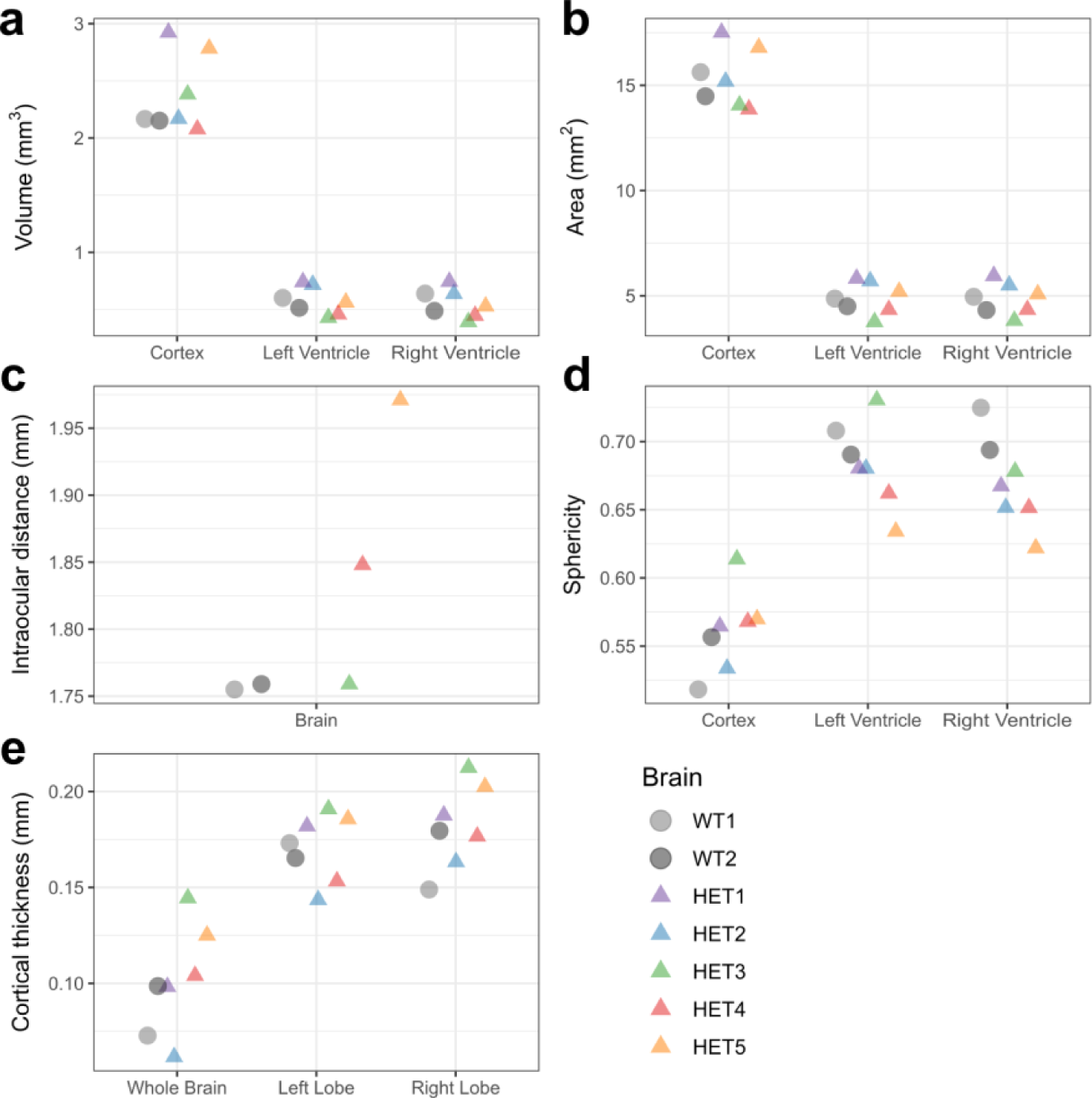
Morphological characterization of E12.5 wild-type and *Chd8*^*+/-*^ mouse brains. **(a-c)** Cortical and ventricular **(a)** volume, **(b)** surface area, and **(c)** intraocular distance. **(d)** Sphericity of cortex and ventricles. **(e)** Cortical thickness of whole brains and left and right lobes. Quantifications were extracted from ventricular and cortical iso-surfaces.

*CHD8* mutant patients often present craniofacial abnormalities (7). We report a slight increase in the intraocular distance and variability within the *Chd8*^*+/-*^ group (wild-type [n=2] 1.75 ± 0.002 mm, *Chd8*^*+/-*^ [n=3] 1.85 ± 0.10 mm) (Fig. 2c), which is consistent with mouse studies from similar genetic backgrounds (10). Moreover, we observed a slight decrease in ventricular sphericity in *Chd8*^*+/-*^ mice (wild-type [n=4] 0.70 ± 0.01, *Chd8*^*+/-*^ [n=10] 0.67 ± 0.03) (Fig. 2d). We also found a higher variability in whole-brain cortical thickness in *Chd8*^*+/-*^ mice (wild-type [n=2] 0.08 ± 0.02 mm, *Chd8*^*+/-*^ [n=5] 0.10 ± 0.03 mm) (Fig. 2e). Thus, only some craniofacial phenotypes were detected during early embryonic development.

### Quantifying apical cell morphology in Chd8^+/-^ mice

To identify cellular and tissue mechanics abnormalities preceding ASD-associated macrocephaly, we used MorphoGraphX to isolate and mesh the apical boundaries of imaged epithelial cell patches in matching regions of the cerebral cortex (18). By taking tissue curvature into account, we segmented a large number of cell outlines from one *Chd8*^*+/-*^ [n=3854] and one wild-type [n=1031] sample (Fig. 3a-b).

**Figure 3.**
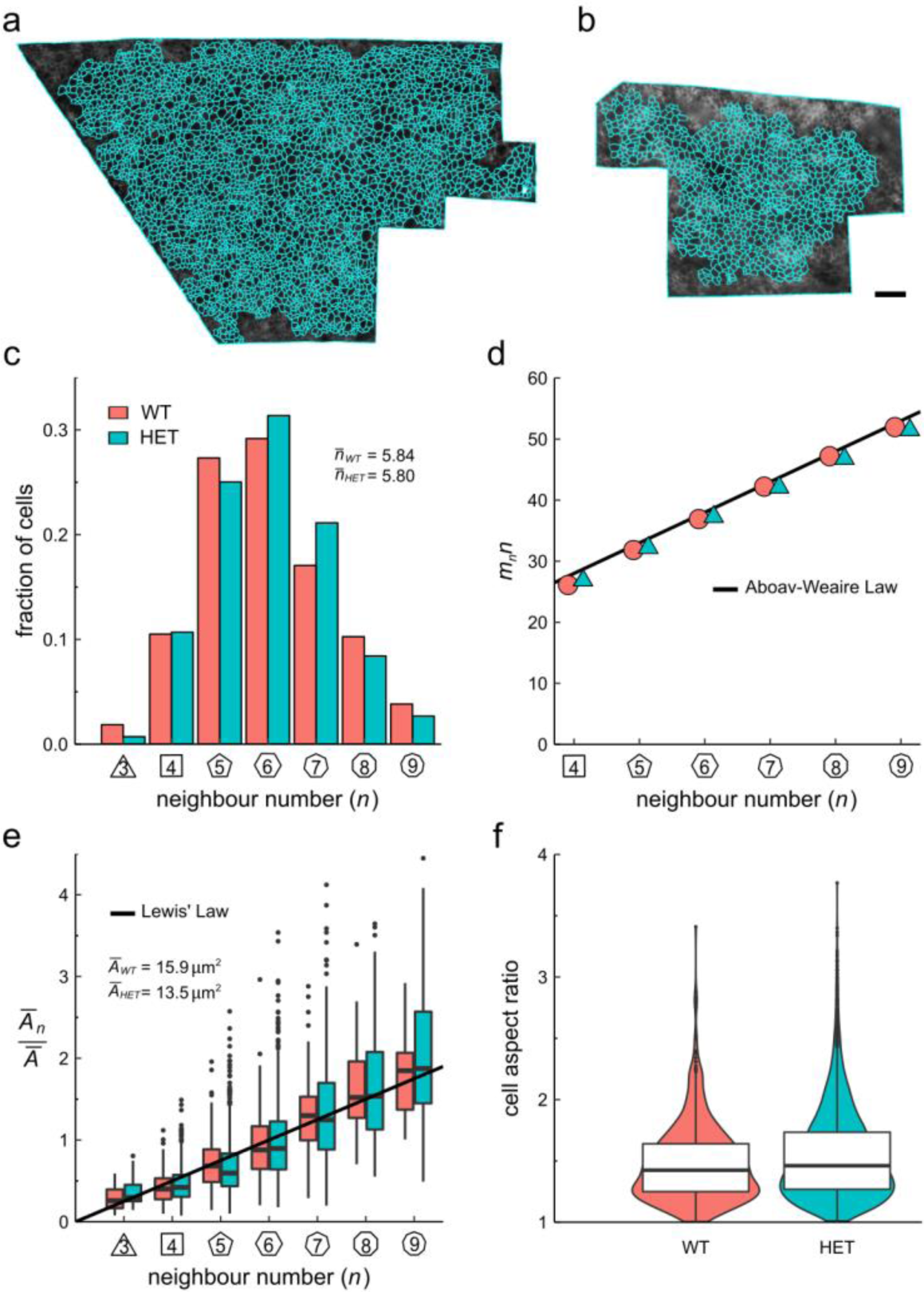
Quantification of apical cell morphology in wild-type [n=1031] and *Chd8*^*+/-*^ [n=3854] E12.5 mouse brains. **(a, b)** 2.5D segmentation overlay of the apical surface of **(a)** *Chd8*^*+/-*^ (HET) and **(b)** wild-type (WT) neurocortical epithelium. Scale bar: 20 µm. **(c)** Distribution of apical neighbour numbers per sample. The average number of cell neighbours is 5.84 for WT and 5.80 for HET, which is close to the topological requirement of 6. **(d)** Polygon type *n* times the mean polygon number of neighbours *m* of the cell *n* follows a linear relationship termed Aboave-Weaire’s Law. **(e)** Average apical cell area by cell neighbour number following a linear relationship termed Lewis’ Law (black line). **(f)** Cellular aspect ratios between their longest and shortest axis.

Apical organization was quantified by geometrical properties such as cell number of neighbours, areas, and aspect ratio. We found similar hexagon and heptagon frequencies for the heterozygous (wild-type 29% hexagons, 16% heptagons, *Chd8*^*+/-*^ 31% hexagons, 20% heptagons) (Fig. 3c). The observed relation between the polygon type of cells *n* and the average polygon type of their neighbours *m*_*n*_ termed Aboav-Weaire’s Law (21,22), recapitulated results in both the *Drosophila* wing disc and chicken neural tube epithelium (23) (Fig. 3d). Similarly, we compared area distributions per polygon type and found no significant difference between groups. The average area per polygon type followed a linear dependency in both samples; a relationship termed Lewis’ Law (19,20) (Fig. 3e). Moreover, considering local apical curvature, the aspect ratio was determined by fitting an ellipse to each cell outline and extracting the major and minor axes. The aspect ratio distribution of the *Chd8*^*+/-*^ cells was minimally wider than the wild type (Fig. 3f).

## Discussion

*Chd8* haploinsufficient mice display various ASD-like phenotypes that parallel the clinical signature of individuals with de-novo *CHD8* mutations (10,11,13,24). Consistent with retrospective patient head circumference data, mouse models for *CHD8* haploinsufficiency suggest a postnatal onset of abnormal head growth (10,11). In this study, we queried the neuroanatomy of *Chd8*^*+/-*^ and litter-matched E12.5 control mice using light-sheet microscopy to determine whether morphological anomalies in brain and cortical cell shape could preindicate ASD-associated macrocephaly.

The results from a number of longitudinal studies of postnatal volumetric brain changes have implicated neuroanatomical abnormalities in cortical thickness, ventricular morphology, cortical overgrowth, and increased cortical surface area in the complex trajectory of brain development in individuals with ASD (6,25–30). Accordingly, our work characterized cortical thickness along with volumetric features to confirm whether analogous morphological alterations were observable in haploinsufficient mouse brains. We note that in this particular instance, no significant discrepancies in cortical thickness, ventricular and cortical volumes, or surface areas between groups could be determined. In line with the heterogeneous nature of ASD, it is reasonable to assume that other brain regions may be affected instead. Notably, we provide experimental evidence of dissimilarities in ventricular sphericity and intraocular distance that mirror known phenotypes in haploinsufficient adult mice (Fig. 2) (10).

Similarly, a number of aberrations at the cellular scale have been reported in ASD during the establishment of cortical microarchitecture (26). Consequently, we sought to ascertain differences in the cortical organization between groups as defined by patterns of cell geometric features measured on the apical surface (Fig. 3). Having quantified epithelial morphology according to the cell area, aspect ratio, neighbour topology, and adherence to empirical laws such as Lewis’ and Aboav-Weaire’s (Fig. 3), our data showed no departure in the cortical organization between haploinsufficient brains and controls, suggesting similar mechanical behaviour (20,22).

In this work, we present a multi-scale assessment of the embryonic neuroanatomical implications of *Chd8* haploinsufficiency in mice. We propose that an increased understanding of the identified organ-level differences may shed light on the etiology of hypertrophic brain growth. What is more, our approach opens exciting avenues to investigate the presence of cellular alterations in other implicated brain regions and phenotypic differences across diverse *Chd8* haploinsufficient mouse models, all of which have a wide range of dosage-specific, dimorphic, and behavioural signatures (24,31).

## Limitations

Underscoring the complexity of autism, our results did not show statistically significant differences in overall morphology with the exception of slight deviations in ventricular sphericity and intraocular distance (Fig. 2). Furthermore, our study did not identify aberrations in cortical cellular architecture between groups (Fig. 3). We acknowledge that as only a small sample size could be studied (wild-type [n=2], and *Chd8*^*+/-*^ [n=5]), small morphological differences may have been missed due to the lack of statistical power.

## Abbreviations

ASD: autism spectrum disorder
CHD8: chromodomain helicase DNA-binding protein 8
LOF: loss-of-function
CUBIC: clear unobstructed brain/body imaging cocktails and computational analysis
WT: wild type
HET: heterozygous

## Declarations

### Ethics approval and consent to participate

All animal experiments conducted in the USA followed Public Health Service (PHS) policy and guidelines on humane care, and the use of laboratory animals was approved by the Massachusetts Institute of Technology Committee for Animal Care (CAC).

### Consent for publication

Not applicable.

### Availability of data and materials

The datasets used and/or analyzed during the current study are available from the corresponding author on reasonable request.

### Competing interests

The authors declare that the research was conducted in the absence of any commercial or financial relationships that could be construed as a potential conflict of interest.

### Funding

This work has been supported through an SNF Sinergia grant to DI.

### Authors’ contributions

The study was designed by DI. Staining was done by OM, clearing and imaging by HG. HG and LH analyzed the data and wrote the manuscript. All authors read and approved the final version of the manuscript.

## Acknowledgments

We would like to thank Randall Platt and Ashwin S. Shetty for providing the embryos. Moreover, we acknowledge Richard S. Smith for his expert advice on extending MorphoGraphX to enable aspect ratio quantifications of segmented cell outlines.

## Notes

### Competing Interest Statement

The authors have declared no competing interest.

